# Pollen contaminated with field-relevant levels of cyhalothrin affects honey bee survival, nutritional physiology, and pollen consumption behavior

**DOI:** 10.1101/025189

**Authors:** Adam G Dolezal, Jimena Carrillo-Tripp, W. Allen Miller, Bryony C. Bonning, Amy L. Toth

## Abstract

Honey bees are exposed to a variety of environmental stressors that impact their health, including nutritional stress, pathogens, and chemicals in the environment. In particular, there has been increasing evidence that sublethal exposure to pesticides can cause subtle, yet important effects on honey bee health and behavior. Here, we add to this body of knowledge by presenting data on bee-collected pollen containing sublethal levels of cyhalothrin, a pyrethroid insecticide, which, when fed to young honey bees, results in significant changes in lifespan, nutritional physiology, and behavior. For the first time, we show that when young, nest-aged bees are presented with pollen containing field-relevant levels of cyhalothrin, they reduce their consumption of contaminated pollen. This indicates that, at least for some chemicals, young bees are able to detect contamination in pollen and change their behavioral response, even if the contamination levels do not prevent foraging honey bees from collecting the contaminated pollen.

## Introduction

Pollinators are a critical element in healthy ecosystems and key players in sustainable crop production (Ashman et al. 2004, Klein et al. 2007, Aizen et al. 2009). Native and managed bees are among the most important pollinators, but in recent years pollinator populations have plummeted. This is of major concern to ecosystem health and to food security worldwide (Gallai et al. 2009), and the repercussions may contribute significantly to global environmental change (Aizen et al. 2009). Pollinator declines are an area of vigorous research, and all indications are that these declines are the result of multiple, interacting stressors including habitat degradation, agricultural intensification, environmental toxins such as insecticides, and an increased spread of bee diseases as a result of globalization (Oldroyd 2007, Naug 2009, Goulson et al. 2015).

Among these factors, there has been increasing concern over the effects of sublethal doses of pesticides on bee health. While it is clear that high levels of insecticides can cause mortality in honey bees (Johnson et al. 2010), many recent studies have shown that lower, sublethal doses can have effects on many facets of honey bee biology, including behavior (Suchail et al. 2001, Williamson et al. 2013), learning (Decourtye et al. 2004, Frost et al. 2013), colony development (Dai et al. 2010, Wu et al. 2011, Gill et al. 2012), and increased susceptibility to several pathogens (Pettis et al. 2012, Pettis et al. 2013, Doublet et al. 2014). Honey bees encounter pesticides, at lethal and sublethal doses, through a variety of routes. For example, whole hives and even apiaries can be exposed to insecticidal sprays or contaminated planter dust and beekeepers often introduce miticides directly into hives. Foraging honey bees can be exposed directly to pesticides on treated plant materials, and hive bees can be exposed through plant products returned to the hive by foragers (Johnson et al. 2010, Krupke et al. 2012). These multiple routes of exposure contribute to the presence of a wide variety of pesticides inside of most honey bee hives (Mullin et al. 2010). It is well-known that pesticide-contaminated pollen is often collected by foraging workers and brought into colonies; exposure to this pollen can then result in negative impacts on bee health (Pettis et al. 2012, Pettis et al. 2013, Doublet et al. 2014).

One common class of insecticides, the pyrethroids, is used on a wide variety of crops, including many orchard crops such as almonds, apples, and cherries (Epstein et al. 2000), and is the most prevalent class of insecticides found in bee-collected pollen (Mullin et al. 2010). Pyrethroids have reported repellant effects on honey bee foragers, and exposure causes already-foraging bees to decrease foraging activity (Fries and Wibran 1987, Rieth and Levin 1988, Decourtye et al. 2004) and increases the number of non-foraging behaviors exhibited by these foragers (Cox and Wilson 1984). Both of these behavioral changes result in an overall reduction of colony foraging activity (Fries and Wibran 1987, Rieth and Levin 1988). Nonetheless, pyrethroids are commonly found inside honey bee hives (Mullin et al. 2010), showing that, even if there is reduced foraging, pyrethroid-contaminated pollen is being collected by forager bees at a non-negligible level. However, while foraging workers collect this pollen and bring it into the hive, they rarely consume or store pollen themselves; instead, younger hive bees accept, process and consume this pollen (Winston 1987). After contaminated pollen reaches the hive, there is still much we do not know about how hive bees accept or reject such pollen and to what extent it is consumed. This is a key gap in our knowledge, and filling it would provide important information about actual exposure of hive bees to insecticides and their health effects, as well as valuable information about whether bees have behavioral mechanisms that allow them to avoid contaminants in their food.

To address these gaps in our knowledge about the responses of bees to field-relevant doses of pesticides, we took advantage of some readily available, bee-collected pollen that was discovered to be contaminated with lambda-cyhalothrin, a common pyrethroid insecticide (e.g., Karate®). Cyhalothrin levels in this pollen were moderate to high, containing levels below the reported LD_50_ (790 ppb), but higher than the average found in previous surveys of bee hives (Mullin et al. 2010). We first performed a series of experiments that tested the effects of bee-collected pollen from several different plant sources, each contaminated with different levels of cyhalothrin, on the survival, nutritional physiology, and pollen consumption behavior of young, laboratory-kept bees. Next, we used a more refined approach by comparing matched pollen sources that had been experimentally spiked with controlled, field-relevant doses of cyhalothrin. We then observed pollen consumption behavior in both cages of bees in the laboratory and in small nucleus hives kept in the field. Our data show effects of pollen source and contamination on survival and nutritional physiology, and also show that young honey bees change their behavior towards pollen contaminated with this insecticide to reduce pollen consumption.

## Methods

### Bee collected pollen acquisition and pesticide testing

We purchased approximately 5 kg of bee-collected, corbicular pollen that had been pooled from hives in a single apiary in southern Minnesota from a commercial, non-migratory beekeeper. All pollen had been collected in a period of less than one week in May 2012. We then sorted the corbicular pollen pieces by color, which is commonly used as a rough metric for species differences (Schmidt et al. 1987). Subsequently, we used molecular methods to identify the major plant species that was the source of each of the sorted pollen types by following a barcode protocol using the chloroplast *rbcL* gene sequence (Little et al. 2004). The complete blend of pollen contained at least 5 pollen species, determined by pollen color, with the most abundant being *Taraxacum sp.* (dandelion) and *Salix sp.* (willow), each of which made up approximately 8% by mass of the total blend. The polyfloral blend, sorted *Taraxacum*, and sorted *Salix* were then sent in 3 g aliquots to the USDA-AMS-NSL in Gastonia, NC for pesticide residue analysis using GC-MS (Lehotay et al. 2005, Mullin et al. 2010). This screening revealed that the polyfloral blend contained 82.5 ppb of cyhalothrin, *Salix* contained 10.6 ppb, and *Taraxacum* contained 280 ppb, all of which are substantially lower than the previously calculated LD_50_ dose for honey bees, 790 ppb (Mullin et al. 2010). Samples were also screened for 173 other pesticides, and only one other was detected - the herbicide atrazine. Atrazine was detected in the polyfloral blend and *Salix* at levels (12 ppb and 17 ppb, respectively), similar to the mean found by Mullin et al. (2010, 13.6 ppb), which are unlikely to cause mortality effects in caged bees (Helmer et al. 2014).

### Feeding of caged bees on unmanipulated bee-collected pollen

In August and September 2012, we performed three replicates of experiments where cages of 60 bees were fed either no pollen (replicate 1=7 cages, replicate 2 = 8 cages, replicate 3=20 cages, total n=35 cages), a polyfloral blend of pollen (replicate 1=7 cages, replicate 2=9 cages, replicate 3=19 cages, total n=35 cages), *Salix sp.* pollen (replicate 1=5 cages, replicate 2=5 cages, replicate 3=5 cages, total n=15 cages) or *Taraxacum sp.* pollen (replicate 1=6 cages, replicate 2=5 cages, replicate 3=11 cages, total n=22 cages). Cages were set up by removing frames from at least three hives in our research apiary, removing adult bees by brushing, and then keeping frames overnight to allow collection of newly emerged bees the following day. Bees used for the experiment were from healthy colonies, and showed no substantial infection with common honey bee viruses (Supplementary information). Next, 60 bees were counted out by hand into the small acrylic cages (10.16 cm × 10.16 cm × 7.62 cm), which were then stored in an incubation room at 32°C and 50% relative humidity. After the addition of bees to all cages, cages received approximately 0.2 grams wet weight of ground, bee-collected pollen (or no pollen) and had *ad libitum* access to a feeder of 50% sucrose solution. Pollen was replaced daily for the first 7 days (pilot experiments showed cessation of pollen consumption by this time), and cages were monitored for mortality daily for 26 days, after which mortality in some treatments was too high to continue. After the first two independent replicates of this design, we anecdotally observed that bees fed the *Taraxacum* pollen pushed pollen out of their cages (Fig. 1). Therefore, in the third replicate, we also recorded pollen consumption in each cage (polyfloral n=20 cages; *Salix* n=6 cages; dandelion n=11 cages). To monitor pollen consumption, we added precisely 0.2 grams of wet weight pollen to each cage daily, and then carefully removed any remaining pollen 24 hours later. This pollen was then dehydrated in a drying oven for 48 hours, and its mass compared to the dry weight equivalent of 0.2 grams wet weight pollen from the same source. In all three replicates, we also collected a subset of 2 bees from each cage at day 14 for analysis of lipid content and the presence of viruses (Supplementary information).

**Fig. 1:**
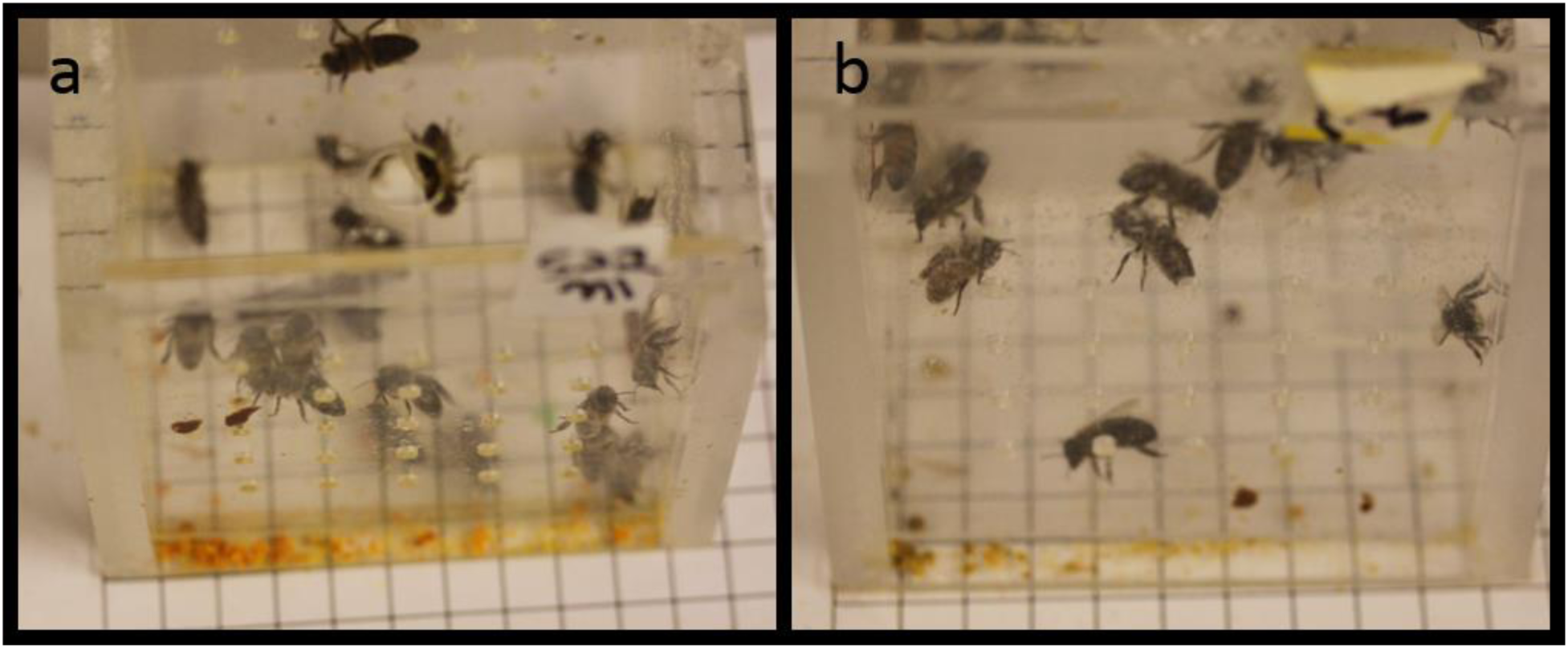
Rejection of cyhalothrin-contaminated pollen. Representative images are shown for a) Cages of bees fed *Taraxacum sp.* pollen that was contaminated with cyhalothrin. After 24 hours, a significant amount of pollen (orange debris at bottom) was not consumed and was pushed under the door of the cage. b) Little pollen remains in cages fed polyfloral or *Salix sp.* because this pollen was not rejected by bees.

To analyze differences in survival between cages in different treatments, we created a survival table and used a Cox proportional hazards regression model using the package [survival] and the R function (coxph), controlling for cage number, variation in starting population, and replicate number. Unless noted otherwise, subsequent statistical analyses were also performed using R. To adjust for multiple comparisons, we used the Benjamini-Hochberg false discovery rate procedure (Benjamini and Hochberg 1995), resulting in an adjusted alpha value of 0.033. For the final replicate, we evaluated differences in pollen consumption by comparing the total pollen consumed per bee during the 6 days of pollen consumption. Since data did not fit normality or homogeneity of variance assumptions, we tested for differences using a Kruskal-Wallis ANOVA followed by a Steel-Dwass posthoc test using JMP statistical software.

### Lipid analysis

After collection, the gut of each bee was removed to prevent food stored in the gut from influencing the results. Bees were then processed for lipid quantification using a phospho-vanillin spectrophotometric assay commonly used on honey bees (Toth and Robinson 2005). Lipid content concentration from two bees per cage inform the different treatment groups was compared, with source cage as a factor, using an ANOVA model followed by a Tukey HSD posthoc test.

### Feeding of caged bees on experimentally contaminated pollen

In June 2014, we mixed newly emerged bees derived from 6 apiary colonies, and then counted 30 bees into each cage, setting up 12 cages per treatment group. Cages received either unmanipulated polyfloral pollen (identical to previous cages, and thus containing 82.5 ppb cyhalothrin), or polyfloral pollen experimentally contaminated with either 140 ppb, 280 ppb, or 560 ppb cyhalothrin. We used the same pollen source that had been used in previous experiments, which had been stored frozen at -20°C. While some degradation to pollen nutritional quality could have occurred, frozen pollen maintains substantial nutritional quality even after years of frozen storage (Dietz and Stevenson 1980). Furthermore, comparisons were only made within experiments using the same-aged pollen, and therefore any quality degradation will not affect our interpretation of the results. These doses were chosen because the field-collected *Taraxacum* pollen contained 280 ppb of cyhalothrin, and hence these doses should represent a range of reasonable field-relevant levels of this insecticide. We assume that no changes in cyhalothrin concentration occurred in our stored pollen stock, as cyhalothrin is stable for multiple years when frozen (Drew et al. 2007). To produce the contaminated pollen, we used laboratory quality lambda-cyhalothrin (ChemService, West Chester, PA, USA), which we diluted into solution in distilled water. Then, 70 μl of water or insecticide solution was added to 0.2g (wet mass) pollen. Cages then received one of the different cyhalothrin treatments daily for 8 days, and mortality was recorded for 14 days, which is where the largest differences in mortality had been observed in the first experiment. Pollen consumption was recorded using the same methods described above. In all treatment groups, very little pollen was consumed on day 1, so this day was removed from analysis. To analyze differences in survival, we used the same Cox proportional hazard method described above. To compare pollen consumption over time, we used pairwise repeated measures ANOVA, controlling for cage and day (time) to prevent overinflation of sample size. To adjust for multiple comparisons between the pairwise tests, we used the Benjamini-Hochberg (Benjamini and Hochberg 1995) false discovery rate procedure, resulting in an adjusted alpha value of 0.0214.

### Feeding of field bees in nucleus hives with experimentally contaminated pollen

To test these effects in a field setting, we created three nucleus hives in our research apiary in October 2014; each hive contained 5 frames of adult bees (approximately equal populations, estimated 8000 bees), one frame of capped brood, one frame containing open brood, and a “pseudo-queen” (queen mandibular pheromone dummy, Mann-Lake, LTD, Hackensack, MN, USA) in lieu of a real queen (Toth and Robinson 2005). All frames used contained little pollen; what pollen was present was removed by scraping out of those cells. Throughout the remainder experiment, forager bees from all treatment groups were free to bring in pollen from outside sources. A 3 cm wooden ring was added to the top of each hive to allow a small dish of pollen to be added on the top bars of the frames without touching the lid of the hive. This approach was repeated three times, with three different nucleus hives, with a total of 9 independent hives over a three week period in October 2014. We prepared polyfloral pollen with no added cyhalothrin, 280 ppb cyhalothrin, or 560 ppb cyhalothrin final concentration, as described above, but scaled to 5 g. Each day, 5 g of the appropriate pollen was weighed into a small plastic dish, which was then placed on the center top bar of a hive. With this arrangement, the pollen dish was inside of the hive and gave hive bees access to the pollen for consumption. Twenty four hours later, the dish was removed, the pollen was dried for 48 hours in drying oven, and the dry mass recorded. To control for hive-level effects, treatments were cycled across nucleus hives each day, so that each nucleus hive received each treatment, and a different treatment each day for 5 days per replicate (with a total of 15 days observed) Due to the effects of weather (rain, cold nights), there was large variation in the amount of pollen consumed each day (i.e., on some days, almost no pollen was consumed in any treatment, as bees remained clustered in the hive). To control for this, we calculated an average amount of pollen consumed among the focal hives each day, and then compared the amount of pollen a hive consumed to that average. This allowed for normalization for days in which very little pollen was consumed across the experiment versus days when a large quantity was consumed. Using the quantity of pollen consumed above the daily average for each hive, we compared pollen using pairwise repeated measures ANOVA controlling for date of observation and hive. To adjust for multiple comparisons, we used Benjamini-Hochberg adjustment of 0.01667.

## Results

### Cages fed unmanipulated field pollen

### Effect of field-collected pollen on survival

First, we determined whether pollen source affects survival in caged bees. We fed caged honey bees field-collected pollen that contained moderate levels of cyhalothrin insecticide contamination. Our screening revealed that polyfloral pollen contained 82.5 ppb, *Salix* contained 10.6 ppb, and *Taraxacum* contained 280 ppb, all of which are lower than reported LD_50_ doses, but higher than previous reports of in-hive contamination of pollen (Mullin et al. 2010). Over a 26 day period (Figure 2), survival of caged bees was not significantly different between bees fed no pollen (n=35), polyfloral pollen (n=35), or *Salix sp.* pollen (n=15) (Cox proportional hazard model, alpha corrected <0.033, p>0.033), but survival was significantly lower in the bees fed the *Taraxacum sp.* pollen (n=22) compared to those fed the polyfloral pollen (p =0.005), *Salix sp.* pollen (p=0.0005), or even bees fed no pollen at all (p=0.014). Overall, there were no significant difference in survival in the bees fed polyfloral, *Salix sp.,* or no pollen, and *Taraxacum sp-* fed bees exhibited lower survival than any of the other groups.

**Fig. 2:**
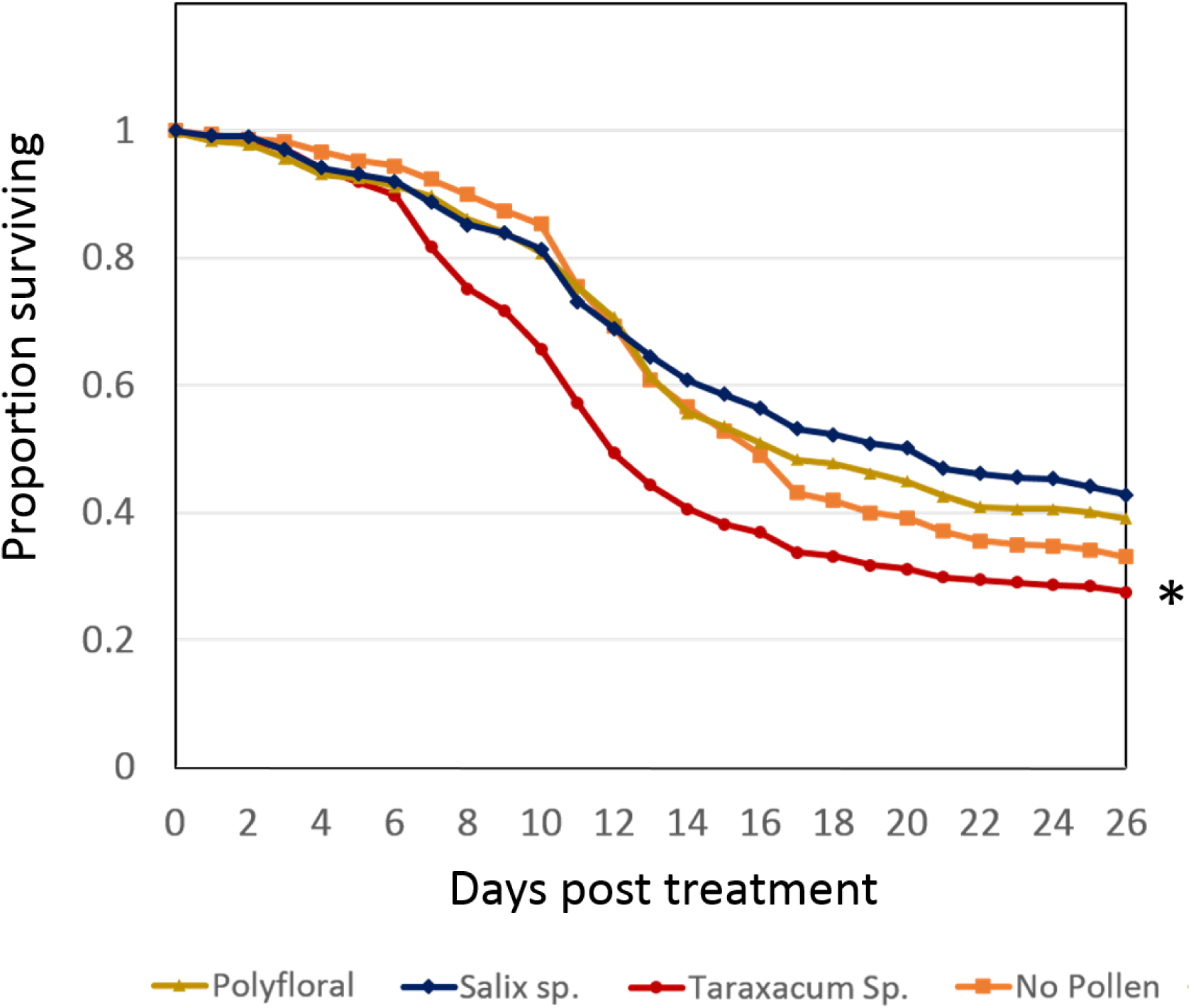
Proportion of original 60 bees surviving per cage averaged across all cages in a treatment. Asterisk indicates significant difference: significantly fewer *Taraxacum*-fed bees survived compared to all other groups, which did not differ from each other (Cox proportional hazards model, alpha adjusted p<0.033).

### Differential consumption of field-collected pollen

To distinguish whether the above effects on honey bee survival were due to consumption of unhealthy pollen or due to reduced pollen consumption (or a combination of the two), we observed consumption levels of the different pollen sources. In cages of honey bees fed field-collected pollen, there were some significant differences in total pollen consumption between the groups (Fig. 1, 3). Over 5 days, the pollen consumed per bee per day did not differ between cages fed polyfloral pollen (n=20) and *Salix sp.* (n=6) (Kruskal-Wallis ANOVA, Steel-Dwass multiple comparison, p>0.05), but bees fed *Taraxacum sp.* pollen (n=11) consumed significantly less pollen than bees in cages fed polyfloral pollen (Kruskal-Wallis ANOVA, Steel-Dwass multiple comparison, p<0.05) or *Salix sp.* pollen (Kruskal-Wallis ANOVA, Steel-Dwass multiple comparison, p<0.05).

**Fig. 3:**
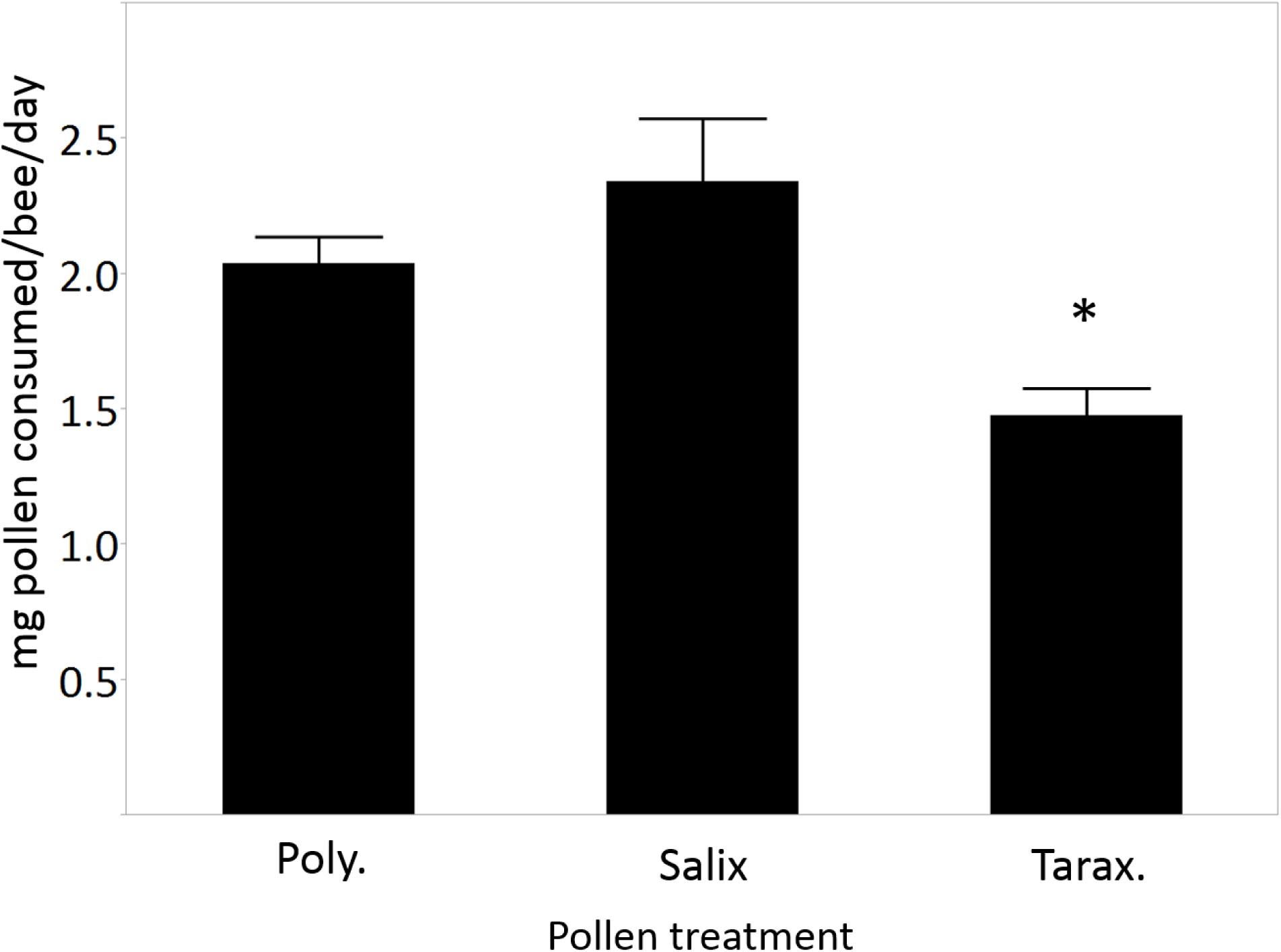
Mean +/-SE milligrams of pollen consumed per bee per day of 6 days in cages fed polyfloral, *Salix sp.,* or *Taraxacum sp.* pollen. Asterisk indicates significant differences: *Taraxacum*-fed bees consumed significantly less pollen than the other groups (Kruskal-Walls ANOVA, Steel-Dwass posthoc test, p<0.05)

### Honey bee lipid content

Lipid content is an indicator of bee health (Wilson-Rich et al. 2008), therefore we measured lipid content of the caged bees fed different pollen diets. As with survival, pollen diet also resulted in significant differences in their whole-body lipid content at day 14 (ANOVA, p<0.05). Polyfloral pollen-fed bees (n=40) did not have significantly different lipid stores than bees fed *Salix sp.* (n=16; Tukey HSD, p>0.05), but did have higher lipid levels than bees fed no pollen (n=40) and bees fed *Taraxacum sp.* pollen (n=19; Tukey HSD, p<0.05). There were no significant differences in lipid content between bees fed *Salix sp.,* no pollen, or *Taraxacum sp.*, showing that, overall, bees fed polyfloral pollen had the highest lipid content, bees fed *Salix sp.* had intermediate lipid stores (not significantly different from any other treatment group) and bees fed no pollen or *Taraxacum sp.* had the lowest (Fig. 4).

**Fig. 4:**
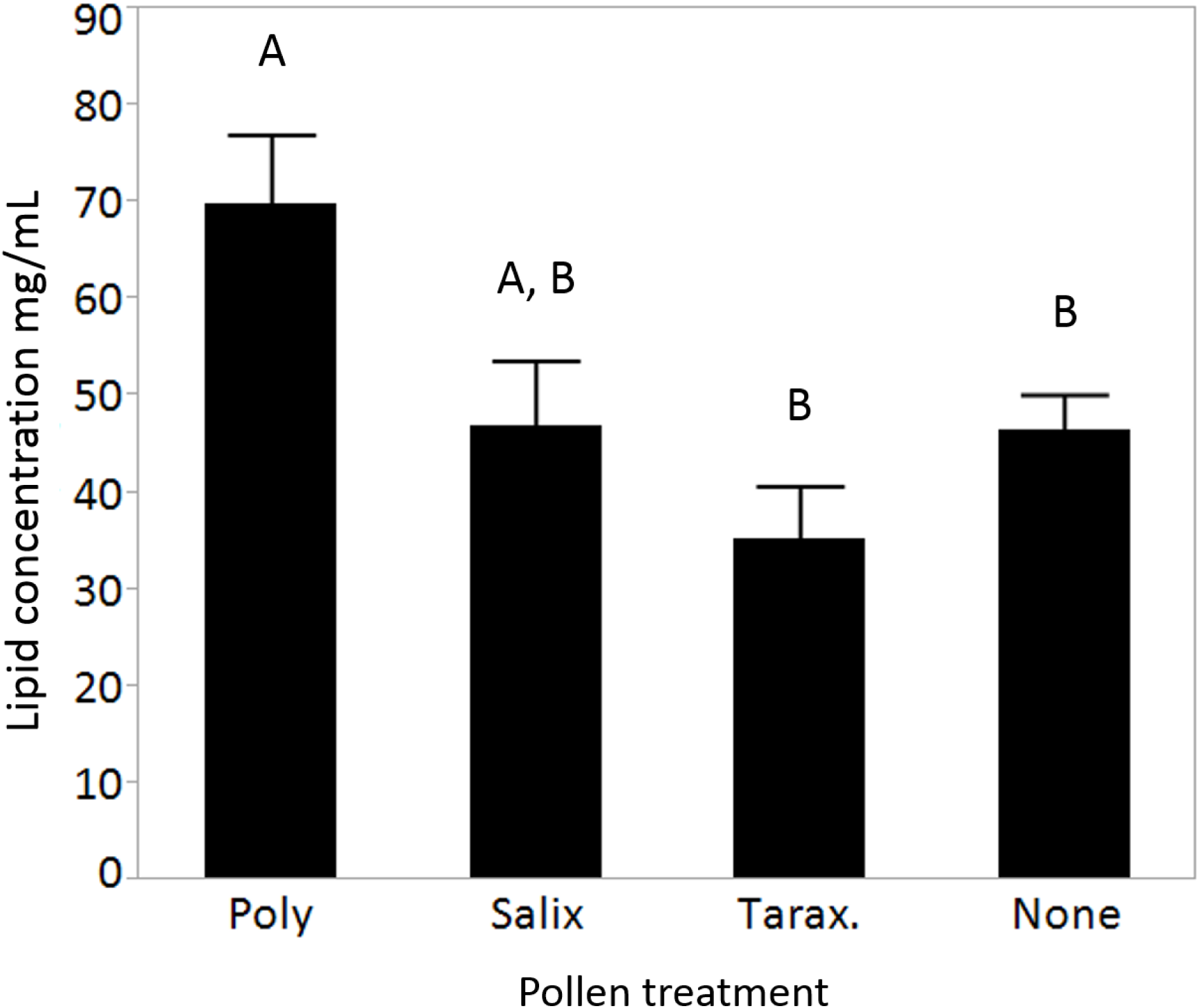
Mean +/- SE lipid concentration (mg/mL) of 14-day-old bees collected from cages fed polyfloral pollen, *Salix sp.* pollen, *Taraxacum sp.* pollen, or no pollen. Letters denote significant differences (Tukey HSD, p<0.05).

### Effect of feeding pollen artificially treated with cyhalothrin insecticide

### Mortality effects of cyhalothrin-treated polyfloral pollen

Based on our findings from cages fed unmanipulated pollen, we hypothesized that the observed effects were caused by the sublethal contamination of some of the pollen sources, e.g. the *Taraxacum sp.* pollen, with cyhalothrin. However, this finding was confounded by the different levels of contamination in different pollen sources. Therefore, we artificially contaminated our polyfloral pollen blend with cyhalothrin levels similar to those found in field-collected pollen. In these cages, there were no significant differences in mortality due to pollen contamination over the 14 day observation period (Cox Proportional Hazard model, p>0.05, Supplemental Fig. 1).

### Pollen consumption

We also evaluated whether pollen consumption was influenced by pollen source or cyhalothrin contamination alone. When pollen consumption in the cages that received cyhalothrin-contaminated polyfloral pollen was compared over time, the same pattern seen in cages fed field-collected pollen was observed (Fig. 5). Bees in control cages consumed significantly more pollen over time than bees fed pollen with 140 ppb cyhalothrin (repeated measures ANOVA, d.f.=1, 5; F=21.44; alpha corrected <0.0214, p=0.0057), 280 ppb cyhalothrin (repeated measures ANOVA, d.f.=1,5; F= 52.62; alpha corrected <0.0214, p=0.00078, n=12) or 560 ppb cyhalothrin (repeated measures ANOVA, d.f.=1,5; F=16.88; alpha corrected <0.0214, p<0.00928).

**Fig. 5:**
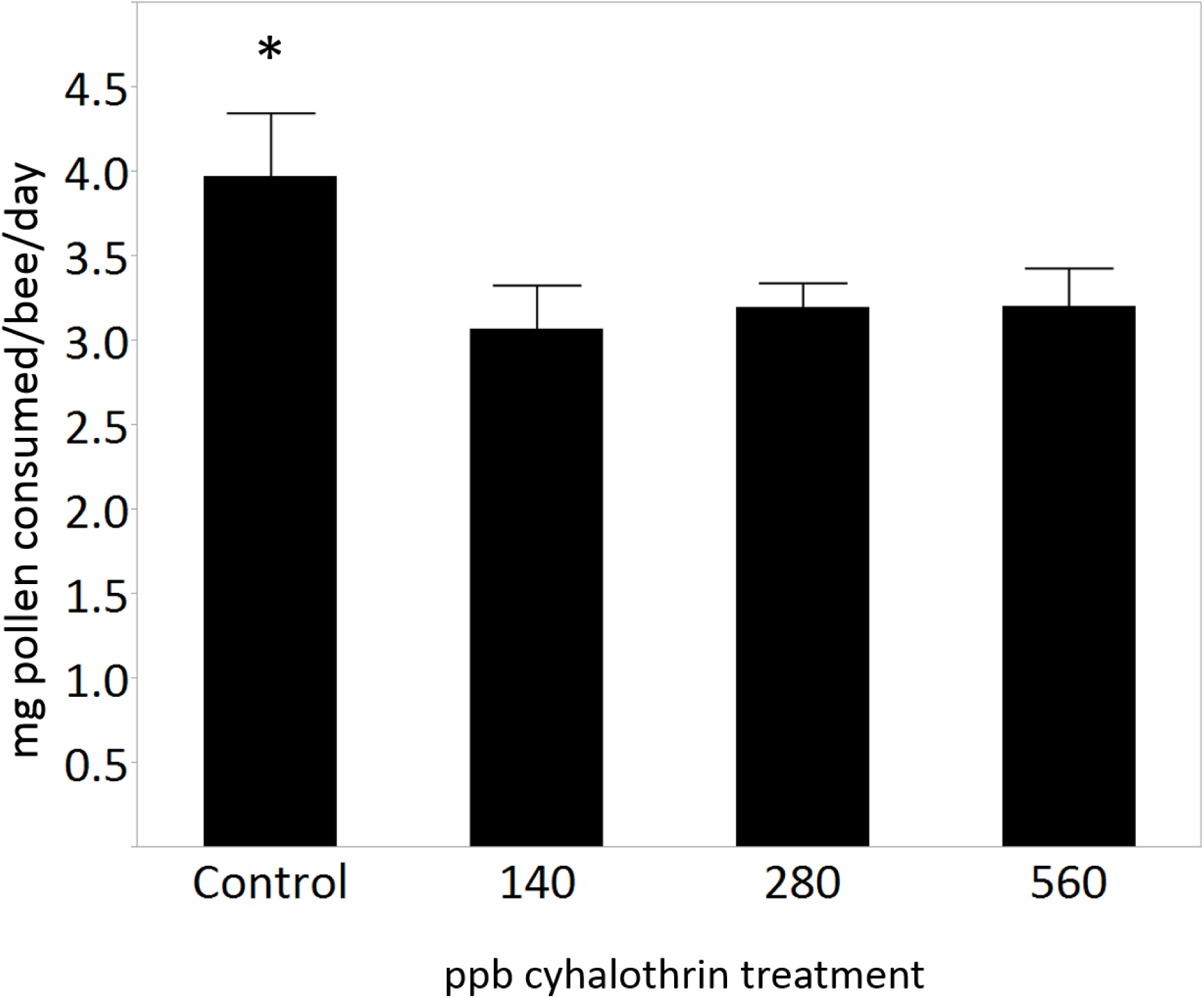
Mean +/- SE milligrams of pollen consumed per bee per day over 7 days in cages fed unmanipulated polyfloral pollen (control) or pollen contaminated with 140, 280, or 560 ppb cyhalothrin; asterisk denotes significant differences (repeated measures ANOVA, p<0.0214)

### Nucleus hives fed pollen artificially treated with cyhalothrin

### Pollen consumption

To evaluate if the differences in pollen consumption observed in cages would occur in more field-relevant conditions, we used small nucleus hives in the field to test honey bee consumption of polyfloral pollen with three different cyhalothrin levels: no added cyhalothrin (note levels are not zero, but measured at 82.5 ppb), 280 ppb cyhalothrin-contaminated, and 560 ppb-contaminated. We found that when nucleus hives were presented with polyfloral pollen containing either no added cyhalothrin, the same dose found in field pollen (280 ppb) or double that dose (560 ppb), the highest dose resulted in significantly reduced consumption of pollen compared to control (Fig. 6). We found that hives consumed significantly less pollen over the daily average when treated with 560 ppb cyhalothrin than controls (repeated measures ANOVA, d.f.=1,9; F=22.65; p<0.01667), but consumption of pollen treated with 280 ppb cyhalothrin was not significantly different between consumption of control pollen and 560ppb cyhalothrin pollen (repeated measures ANOVA, d.f.=1,9; F=1.57; p>0.05).

**Fig. 6:**
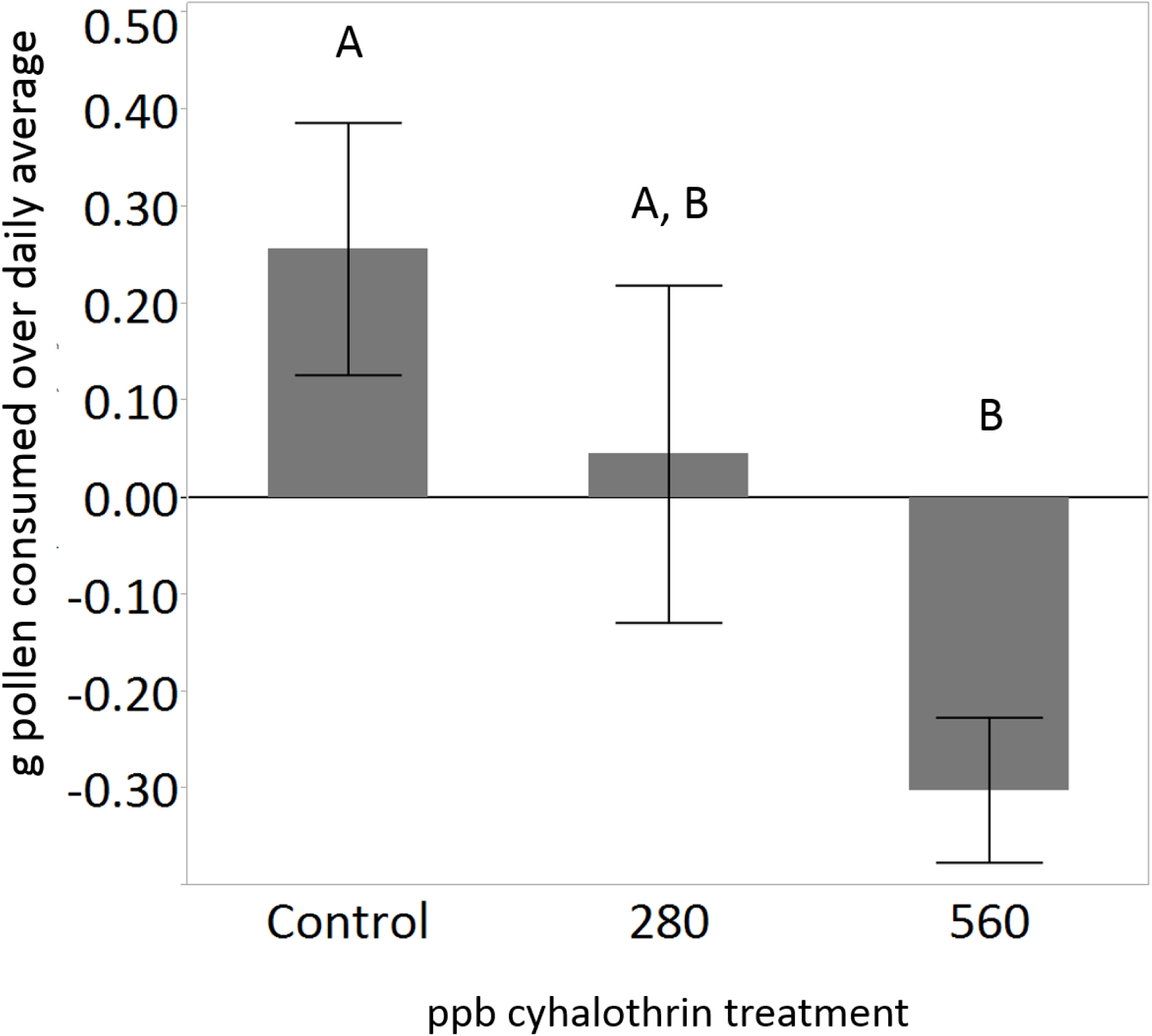
Mean +/- SE of the proportion of pollen consumed over the daily average when nucleus hives were presented with unmanipulated polyfloral pollen (control) or polyfloral pollen treated with 280 ppb or 560 ppb cyhalothrin. Letters denote significant differences (repeated measures ANOVA, p<0.01667).

## Discussion

Here, we present new data adding to the large body of literature showing that exposure to pesticides in the environment can have important sublethal effects on honey bees that may not be evident in short term monitoring for lethal exposure (Suchail et al. 2001, Decourtye et al. 2004, Dai et al. 2010, Wu et al. 2011, Gill et al. 2012, Pettis et al. 2012, Frost et al. 2013, Pettis et al. 2013, Williamson et al. 2013, Doublet et al. 2014). We show that exposure to some pollen sources contaminated with cyhalothrin, a common pyrethroid insecticide, can significantly affect long term lifespan in caged laboratory bees and reduce lipid stores. Low lipid stores are associated with compromised nutrition and immunity (Alaux et al. 2010) and may thus have possible consequences to bee health. Furthermore, to the best of our knowledge, we show for the first time that honey bees in cages or hives are capable of detecting sub-lethal levels and reducing pollen consumption when presented with pollen contaminated with cyhalothrin, either directly from the field or using experimentally-treated pollen.

Caged, newly-emerged bees fed on *Taraxacum sp.* pollen that was contaminated with cyhalothrin exhibited significantly lower survival over a 26 day period than bees fed *Salix sp.* or a polyfloral blend, or even bees fed no pollen at all (Fig. 2). While it is well-established that bees survive best on polyfloral pollen, and that individual pollen varieties have different effects on lifespan and pathogen resistance (Schmidt 1984, Schmidt et al. 1987, Di Pasquale et al. 2013), it is surprising that bees fed no pollen at all would survive more than bees fed *Taraxacum sp.* pollen. While *Taraxacum* pollen is not an ideal source for bees, and honey bees are unable to raise brood successfully on a *Taraxacum* diet alone (Herbert et al. 1970), *Taraxacum* pollen has been reported to improve survival versus pollen-less controls, albeit not as much as many other pollen varieties (Schmidt et al. 1987). In any case, as bees commonly forage on *Taraxacum* pollen (Mayer et al. 1991), it could provide a significant source of hive nutrition, especially during the times of year when *Taraxacum* is one of the few plants blooming. Our finding that bees fed on *Taraxacum* pollen survived worse than bees fed on any other treatment groups makes sense in light of the higher cyhalothrin levels detected in this pollen.

We also found that bees fed *Taraxacum sp.* pollen consumed significantly less pollen than counterparts fed polyfloral or *Salix sp.* pollen (Fig. 1 and 4). Because bees will readily forage for and consume *Taraxacum* pollen (Schmidt et al. 1987, Mayer et al. 1991, Alaux et al. 2010), we hypothesized that the change in consumption was due to cyhalothrin contamination. Cyhalothrin has documented repellant properties on honey bees foraging in the field, causing a reduction in foraging behavior (Cox and Wilson 1984, Fries and Wibran 1987, Rieth and Levin 1988, Decourtye et al. 2004); however, our pollen was collected by foraging honey bees in the field, and therefore must have been at a contamination level below a threshold for repellency. Our observations showed, however, that when just young bees (not old bees, such as the foragers that collected the pollen) are presented with this pollen, as they would be in a natural hive setting, they are able to detect contaminants in the pollen and respond by reducing pollen consumption. This reduced consumption was insufficient, however, to fully prevent the detrimental effects of the insecticide (Figs. 2 and 3).

However, it is problematic to make strong inferences about cyhalothrin effects based on observations of bees consuming cyhalothrin in the context of different pollen species. Therefore, we artificially treated our polyfloral pollen blend with cyhalothrin at three concentrations similar to that found in field-collected *Taraxacum sp.* pollen. When caged bees were fed this pollen, bees consumed significantly less pollen, whether the pollen contained 140 ppb (half of field collected), 280 ppb (identical to field collected) or 560 ppb (double field-collected), compared to controls fed pollen without added pesticide contamination (Fig. 5). This shows that, even comparing identical pollen sources, young (nest-aged) bees are less likely to consume pollen containing cyhalothrin at a level acceptable for collection by foraging bees. Surprisingly, when bees were fed polyfloral pollen experimentally treated with insecticide, we did not observe any significant differences in survival over the experimental period. However, since honey bee response to nutrition may buffer them against the effects of pesticides (Schmehl et al. 2014), it is possible that our use of a higher-quality polyfloral diet prevented the subtle change in survival that was observed in bees fed only the *Taraxacum sp.* pollen source. Another possibility is that other environmental differences in the bees, due to differences in year and season of the experiment, contributed to a higher resistance to pesticide effects. Because these experiments were performed on different colonies, genetic variation in pesticide resistance could have affected our results.

Our observation that honey bees consume less pesticide-contaminated pollen was also validated in experiments using small nucleus hives in the field (Fig. 6). Although the difference in pollen consumption was not as strong as the findings from the cages, the trend of high consumption in controls, intermediate in 280 ppb-fed bees, and low consumption in 560 ppb-fed bees fits the same pattern. This is particularly notable because these nucleus hives contained a mixed age demography, compared to cages kept in the laboratory, which is more representative of full-sized hives. Therefore, our findings suggest that, even if foraging honey bees bring cyhalothrin-contaminated pollen into the hive (Mullin et al. 2010), the young bees that actually consume pollen may be less likely to consume cyhalothrin-contaminated pollen. However, more comprehensive choice tests would be necessary to fully test this hypothesis and to determine if the mechanism of detection requires consumption and if exposed bees discriminate only contaminated pollen or exhibit a more generalized rejection of all pollen due to insecticide poisoning.

Lipid content is a common metric for bee health, as it can provide a physiological measurement of diet quality and immunocompetence (Wilson-Rich et al. 2008), and lipid content can have important effects on pathogen resistance (Alaux et al. 2010) and behavior (Toth and Robinson 2005). Here, we found that bees fed polyfloral pollen had, unsurprisingly, higher lipid content than bees fed no pollen, with bees fed a single source of *Salix sp.* showing somewhat lower, but not significantly different lipid levels. It is particularly notable that *Taraxacum sp.-*fed bees exhibited lipid content most similar to those fed no pollen at all. In a previous study by Alaux et al. (2010), bees fed any of multiple pollen sources, including *Taraxacum*, exhibited higher fat stores than those fed no pollen. While it is possible that our *Taraxacum-*fed bees contained lower lipid levels due to their overall reduction in pollen consumption, it seems unlikely that a reduction in pollen consumption would result in lipid levels as low as bees fed no pollen at all, especially knowing that *Taraxacum* normally increases fat stores compared to unfed controls (Alaux et al. 2010). Therefore, the effects on lipid content were likely driven by cyhalothrin contamination in the pollen, suggesting that pesticide exposure could negatively affect nutritional physiology. Furthermore, recent evidence has shown that diet can affect pesticide resistance in honey bees (Schmehl et al. 2014). Together, these findings show how this type of scenario could set the stage for a “vicious cycle” of pesticide contamination, causing damaging effects on nutritional physiology which in turns decreases pesticide resistance, potentially resulting in higher mortality (as observed in our first experiment). Because young bees need to consume pollen to produce hypopharyngeal gland secretions to feed developing larvae (Winston 1987), a reduction in pollen consumption and nutritional physiology in young bees could have trans-generational effects.

Overall, we provide data underlining the increasingly well-established fact that sublethal levels of pesticides are able to enter honey bee hives and can cause subtle physiological and behavioral effects that may be difficult to detect (Suchail et al. 2001, Wu et al. 2011, Pettis et al. 2012, Pettis et al. 2013, Helmer et al. 2014). Furthermore, we present novel data showing that young honey bees, as well as small hives, can detect some types of contamination in pollen and reduce consumption of that pollen. This effect is particularly important because it indicates that, even if contaminated materials are collected by foraging bees, the nest bees may not accept or consume all contaminated pollen. While we have observed this effect due to cyhalothrin contamination, it is not clear if this effect is widespread; it may be due to the reported repellant properties of pyrethroid insecticides (Cox and Wilson 1984, Rieth and Levin 1988). In fact, honey bees and bumble bees actually consume more sugar solution when it contains neonicotinoid insecticide contamination (Kessler et al. 2015); therefore, it is important to investigate how honey bees respond to different chemical contaminants, as they can result in completely opposite responses by bees. While previous reports indicated that pyrethroid insecticides caused external irritation to bees (i.e., the proboscis and antennae (Rieth and Levin 1988)) consumption or tasting of the contaminant may be also be an important component of repellency, especially at lower doses. At lower contamination levels, foragers, which do not consume the pollen they collect, are not repelled by the presence of cyhalothrin, but hive bees that consume the pollen are. Future work will be necessary to investigate if foraging bees and hive bees simply have different thresholds for pyrethroid repellency, or if the ability of nest honey bees to reject pollen contaminated by insecticides is a more widespread mechanism.

## Acknowledgments

We would like to thank Griffin Smith, Arvin Foell, Giselle Narvaez, and Catie Steinfadt for help with the experiments, and Ali Berens and Jennifer Jandt for comments on the manuscript and statistical advice. This work was supported by USDA-AFRI #2011-04894.

